# Neural Tracking of Speech Envelope as an Index of Spatial Release from Masking

**DOI:** 10.64898/2026.06.29.734758

**Authors:** Juan-Daniel Galeano-Otálvaro, Benajmin Dieudonné, Tom Francart, Jan Wouters

## Abstract

Understanding speech in noisy environments relies strongly on binaural cues such as interaural time and level differences (ITDs and ILDs), which support spatial hearing and the segregation of competing sound sources. When these cues are degraded, listeners experience substantial difficulty in complex acoustic environments.

Behavioural measures of binaural benefit, such as binaural masking level differences, binaural intelligibility level differences, and spatial release from masking (SRM), are well established in normal-hearing (NH) listeners, but they require an active behavioural response. Neural speech tracking using electroencephalography (EEG) has emerged as a promising approach for quantifying neural processing of continuous speech, yet its sensitivity to spatial hearing cues remains insufficiently characterised.

We investigated the neural correlates of spatial release from masking in NH listeners using EEG-based neural speech tracking. Nineteen participants listened to continuous Dutch speech stories presented with masking noise under two spatial configurations, collocated (S0N0) and spatially separated (S0N90), across multiple signal-to-noise ratios (SNRs). Neural tracking of the speech envelope was quantified using both envelope reconstruction and temporal response function (TRF) analyses.

Spatial separation enhanced neural tracking of the target speech envelope, particularly at challenging SNRs where behavioural SRM was also observed. TRF analysis further revealed increased amplitudes and decreased latencies of late cortical components consistent with spatial unmasking effects.

These findings demonstrate that neural speech tracking captures cortical signatures of spatial unmasking and closely reflects behavioural improvements in speech understanding. Establishing these relationships in NH listeners supports the development of objective neural measures for evaluating binaural benefit in difficult-to-test populations.

## Introduction

Spatial hearing—the ability to perceive, localize, and segregate sounds based on their spatial origin—is a fundamental function of the auditory system. It relies on the neural processing of binaural cues such as interaural time differences (ITDs) and interaural level differences (ILDs), together with monaural spectral cues shaped by the head and pinnae (Blauert, 1997). Through the integration of these cues, spatial hearing supports the formation of auditory objects and enables the parsing of complex acoustic scenes, particularly in natural, multi-source listening environments (Blauert, 1997; Litovsky et al., 2021).

A central functional role of spatial hearing lies in its contribution to speech understanding in everyday listening conditions. Speech communication rarely occurs in quiet; instead, listeners must typically extract a target talker from background noise or competing speakers. Spatial separation between sound sources allows the auditory system to exploit binaural unmasking and spatial release from masking, thereby improving access to the target speech signal and facilitating selective auditory attention (Dieudonné & Francart, 2019; Litovsky et al., 2021; Plomp, 1976). These benefits reflect not only peripheral acoustic advantages but also central and cortical mechanisms that integrate spatial and linguistic information over time (Li et al., 2024; Zobel et al., 2022).

The behavioural advantages of spatial hearing for speech perception are well established. Improved speech intelligibility with spatially separated sources has been demonstrated across a wide range of acoustic conditions, including varying signal-to-noise ratios (SNRs) and masker types (Best et al., 2012; Biberger & Ewert, 2019; Blauert, 1997; Litovsky et al., 2021). At the neural level, electrophysiological studies have shown that cortical activity more strongly tracks attended than unattended speech, particularly when spatial cues support perceptual segregation (Das et al., 2018; Hausfeld et al., 2020; Kerlin et al., 2010). Together, these findings point to a close relationship between binaural hearing, selective attention, and cortical speech processing.

In parallel, electrophysiological approaches have been used to investigate neural sensitivity to binaural cues, employing event-related potentials (ERPs), frequency-following responses (FFRs), auditory steady-state responses (ASSRs), and auditory change complexes (ACCs). These studies have targeted specific mechanisms such as interaural phase differences, binaural interaction components, and envelope-based cues (e.g., Clinard et al., 2016; Fan & Gifford, 2024; Gransier et al., 2017; Haywood et al., 2015; Laumen et al., 2016; McAlpine et al., 2016; Undurraga et al., 2016; Van Eeckhoutte et al., 2018; Vercammen et al., 2017). While these electrophysiological approaches provide valuable insight into binaural cue encoding under controlled conditions, they often rely on simplified stimuli and probe isolated cue processing rather than continuous, ecologically valid speech perception. Establishing how these electrophysiological measures reflect binaural processing in normal-hearing listeners is therefore a necessary first step toward extending them to continuous, ecologically valid speech and, ultimately, to populations that cannot be tested behaviourally.

More recently, neural speech tracking paradigms have emerged as a powerful tool for studying auditory processing under naturalistic listening conditions. Using electroencephalography (EEG) combined with system identification techniques such as temporal response functions (TRFs), the brain can be modelled as a linear or quasi-linear system that transforms continuous speech features—most commonly the speech envelope—into time-locked neural responses (Crosse et al., 2016, 2021). Both encoding (forward/TRF) and decoding (backward/envelope reconstruction) approaches have demonstrated that neural tracking strength correlates with speech intelligibility (Vanthornhout et al., 2018), attentional focus and listening effort (Decruy et al., 2020; He et al., 2025; Lesenfants & Francart, 2020), and can be robustly measured in cochlear implant users as well (Somers, Verschueren, et al., 2018; Verschueren et al., 2019). Importantly, neural speech tracking has also begun to be applied to research spatial and binaural hearing. Previous work has reported spatial modulation of TRFs and envelope reconstruction accuracy in multi-talker environments (Bednar et al., 2017; Bednar & Lalor, 2018), speech-in-noise paradigms with manipulated binaural cues (Dieudonné et al., 2024), and extensions across developmental populations and competing-speech conditions (Ahmed et al., 2026; Song et al., 2020). These studies, however, have not specifically examined how spatial separation and signal-to-noise ratio relate to neural speech tracking of continuous, natural speech in normal-hearing listeners.

Characterising this relationship is an important step toward a mechanistic understanding of spatial hearing at the cortical level and toward objective neural markers grounded in normal auditory processing. In the present study, we therefore focus exclusively on normal-hearing listeners and investigate how spatial hearing manipulations—specifically spatial separation and variations in signal-to-noise ratio—affect EEG-based neural tracking of continuous, natural speech. By studying these factors in a normal auditory system, this work takes a step toward a neurophysiological reference framework for interpreting spatial hearing effects in more complex or clinical populations.

Signal-to-noise ratio was systematically varied across spatial listening conditions. We hypothesised that spatially driven enhancements in neural speech tracking would be most pronounced at challenging SNRs. Under these conditions, spatial separation is expected to yield measurable increases in envelope reconstruction accuracy, reflecting spatial release from masking at the cortical level. In addition, we (specifically) hypothesised that spatial unmasking would be reflected in the morphology of the TRFs, with increased amplitudes and decreased latencies for spatially separated compared to collocated conditions, particularly at low SNRs. This prediction is motivated by previous auditory evoked potential and envelope tracking studies showing that reductions in masking are associated with increased response amplitudes and decreased latencies, reflecting more robust and temporally resolved cortical representations of the target signal (Dieudonné et al., 2024; Song et al., 2020; Van Hirtum, Somers, Dieudonné, et al., 2023; Van Hirtum, Somers, Verschueren, et al., 2023; Zobel et al., 2022). In contrast, at higher SNRs—where speech is already well represented acoustically—spatial configuration is expected to exert reduced or negligible effects on neural tracking metrics, potentially reflecting ceiling effects or diminished reliance on binaural cues when acoustic clarity is high (He et al., 2025; Rennies & Kidd, 2018).

## Methods

### Participants

Nineteen adults (17 female, median age 20 years, min = 19, max=33) participated in the study. All participants were native Dutch speakers with pure-tone hearing thresholds at or below 20 dB HL at all octave audiometric frequencies from 250 to 8000 Hz. This study was approved by the Medical Ethics Committee Research UZ/KU Leuven (project number S70169). All participants gave written informed consent for their participation in the study.

### Experimental set-up

The experiment took place in a double-walled soundproof room. Stimuli were presented through Genelec loudspeakers (Genelec Oy, Iisalmi, Finland) positioned approximately one metre away from the centre of the head of the participant at 0° and 90° or −90° azimuth. Stimuli were presented via an RME Fireface UCX soundcard (Audio AG, Haimhausen, Germany), using the software platform APEX 4 (Francart et al., 2008). We recorded EEG using a BioSemi ActiveTwo system (Amsterdam, The Netherlands), with 64 active electrodes, at a sampling frequency of 8192 Hz.

### Stimuli

Target speech was always presented from a loudspeaker located in front of the participant (0° azimuth). Stationary speech-weighted noise was presented either from the same frontal loudspeaker (0° azimuth, referred to as collocated) or from a lateral loudspeaker (90° or −90° azimuth, referred to as separated), creating collocated and spatially separated masking conditions. The position of the masker in the separated condition was kept constant for the duration of the session. The masking noise spectrum was derived from the long-term average spectrum (LTASS) of the speech material. Target speech was presented at 55 dB A, and the signal-to-noise ratio (SNR) was manipulated by adjusting the masker level.

### Procedure

Participants completed two parts. In the first one, purely behavioural, they completed an adaptive speech-in-noise procedure using the Flemish Matrix Sentence Test (Luts et al., 2014). The test consists of lists of 20 sentences spoken by a female speaker. Each sentence lasts approximately 2 s and follows a fixed grammatical structure (name, verb, numeral, adjective, object). The adaptive procedure started at −4 dB SNR and after each sentence the participants selected the words they heard from a matrix displayed on a screen in front of them. The SNR was adapted based on the number of correctly identified words according to the procedure described by Brand and Kollmeier (2002).

In the second part participants listened to fragments of stories in Dutch while we recorded EEG responses. Fragments were presented in trials lasting 2.5–5 min. SNR was set to −13, −10, −8, −6, −4, and −1 dB and remained constant throughout each trial. These levels were determined in pilot testing to capture a wide range of performance while avoiding ceiling and floor effects. Trials were presented in pairs, with both trials in a pair sharing the same SNR. Within each pair, the order of the two spatial conditions (separated and collocated) was randomised. The sequence in which the six SNRs were presented was likewise randomised. In total, each participant listened to 5–7.5 minutes of material for each unique combination of SNR and speaker location. Three participants completed single trials of approximately 5 min per SNR and spatial condition (≈60 min EEG total). Two participants completed two trials of 2.5 min each per SNR and spatial condition (≈60 min EEG total). The remaining 13 participants completed three trials of 2.5 min each (≈90 min EEG total). Due to technical reasons data collection failed for one participant at −1 dB SNR; therefore, data for this condition are reported for n = 18, whereas all other SNR conditions include data from n = 19 participants.

### Behavioural Measures

Speech reception thresholds (SRT) were obtained using the Flemish Matrix Sentence Test (Luts et al., 2014). The SRT corresponded to the signal-to-noise ratio (SNR) at which participants correctly identified 50% of the target words and was estimated from the final SNR of the adaptive track (Brand & Kollmeier, 2002). Spatial release from masking was quantified as the difference in SRT (in dB) between the separated and collocated masking conditions. To complement the behavioural SRT measure, subjective speech understanding was assessed after each story trial. Participants were asked to estimate the percentage of the story they understood by answering the question: “How many of the words did you understand (in percent)?”

### EEG Preprocessing and Envelope Extraction

EEG signals were digitized at 8192 Hz and initially high-pass filtered at 0.1 Hz. The data were then downsampled to 512 Hz. Eyeblinks and other artifacts were automatically removed using a Multi-Channel Wiener Filter (Somers, Francart, et al., 2018). Subsequently, the EEG signal was further downsampled to 128 Hz to reduce computational load.

Channels were band-pass filtered between 0.5 and 8 Hz using a linear-phase FIR filter, implemented as separately designed high- and low-pass filters. Each filter was designed with the least-squares method to achieve a stopband attenuation of at least 10 dB and a stopband ripple below 0.25 dB, and the resulting group delay was compensated for so that no phase distortion was introduced. Signals were subsequently re-referenced to the average of all channels and normalised so that each channel had zero mean and unit standard deviation. Trials corresponding to the same SNR × spatial location combination were concatenated for further analysis.

The speech stimuli were processed through a 28-channel gammatone filter bank with equivalent rectangular bandwidth spacing. The envelope of each frequency band was obtained by taking its absolute value and then compressing it using a 0.6 power law (Biesmans et al., 2015). Like the EEG, the envelopes were band-pass filtered between 0.5 and 8 Hz, normalised, and trials corresponding to the same SNR × spatial location combination were concatenated. The resulting signals were then normalised for subsequent neural tracking analyses.

### Backward model

In the neural tracking framework, the stimulus feature—here the speech envelope—is modelled as a linear function of the EEG. Neural tracking of the speech envelope was quantified by computing the Spearman correlation between the original acoustic envelope *s*(*t*) and a reconstructed envelope *ŝ*(*t*) derived from the EEG recorded during stimulus presentation (Crosse et al., 2016, 2021).

The reconstructed envelope *ŝ*(*t*) was obtained by linearly combining the EEG responses *r*(*t*, *n*) measured at time *t* and channel *n*, using a linear decoder *g*(*n*, τ). Because neural responses are not instantaneous, EEG data were included over a post-stimulus integration window of latencies τ. The reconstruction is therefore defined as:

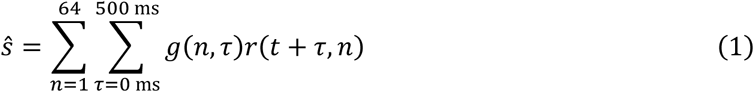

In matrix notation, this can be written as:

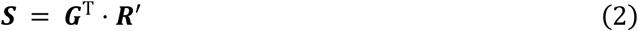

where *R*^′^ represents the matrix of EEG data expanded over channels and time-lagged versions within the integration window, *S* is the speech envelope, and *G* is the decoder vector with dimensions (channels×latencies). The linear decoder thus acts as a spatio-temporal filter that combines information across EEG channels and their delayed responses to optimally reconstruct the speech envelope.

The decoder coefficients were estimated using ridge regression on a training dataset:

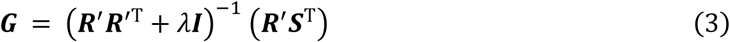

where *λ* is the regularization parameter, set to the maximum absolute value of the auto-correlation matrix ***R***^′^***R***^′T^, and *I* denotes the identity matrix.

For each participant and each listening condition (defined by SNR and spatial location), a condition-specific linear decoder was computed and evaluated using cross-validation. EEG and stimulus data corresponding to each condition were concatenated and divided into 20-seconds folds. For each fold, the speech envelope was reconstructed from the EEG using a decoder trained on all remaining folds. The reconstructed envelopes from all folds were then concatenated to obtain a single reconstructed envelope ŝ(t) per condition, which was subsequently used for neural tracking analysis.

### Forward model

The TRF describes the linear mapping between the stimulus envelope *s*(*t*) and the EEG response *r*(*t*, *n*) at electrode *n*, across a range of temporal delays *τ*. To also include the pre-response integration window, we evaluated lags between -100 to 600 ms. The predicted EEG signal *r̂*(*t*, *n*) can be expressed as the convolution of the stimulus envelope with the TRF weights:

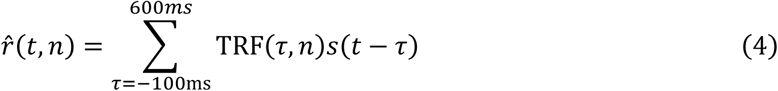

or in matrix notation (for each electrode *n* separately):

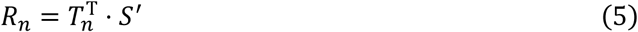

with *S*^′^ as the matrix of the time-lagged versions of the stimulus envelope with dimensions (latencies, time), *R_n_* the EEG signal at electrode *n* with dimensions (1, time), and *T_n_* the TRF coefficients (latencies, 1). The TRF weights were estimated using a regularized regression, as described by Crosse et al. (2016):

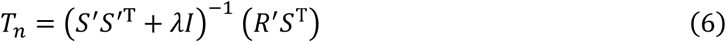

Where λ is the regularization parameter. Here it was set to the maximum absolute value of the autocorrelation matrix *S*^′^*S*^′T^ of the time-shifted stimulus envelope.

For the forward model, we combined the data in the way described for the backward model. Consequently, one TRF was computed per participant, condition, and electrode. Because the analysis focused on the TRF weights themselves rather than reconstruction accuracy, all available data for a given condition were used to estimate the model, no cross-validation procedure was applied.

After calculating the TRFs for each electrode (per participant, per condition), we applied a spatial filter to match the polarity of the TRFs over electrodes following the method applied by Dieudonné et al. (2024). The filter was estimated in two steps. First, reconstruction accuracy was computed per electrode in a cross-validated way, independently for each subject and condition. Second, to estimate a single, common spatial filter, we identified the electrode with the highest average reconstruction accuracy across all subjects and conditions. For each remaining electrode, we computed the correlation between its average TRF and the TRF of this reference electrode. If this correlation was negative, the sign of that electrode’s TRFs was flipped for all subjects and all conditions.

The resulting filter, with coefficients of +1 or −1, was then applied uniformly across all subjects and conditions by averaging over all polarity-matched electrodes, yielding one TRF per subject per condition.

### Peak analysis

Latencies showing consistent effects across subjects were identified using cluster-based permutation testing. At each time point, TRF values across subjects were tested against zero using a cluster-based permutation test (5000 permutations) with sign-flipping across subjects. Consecutive samples exceeding a cluster-forming threshold (cluster α = 0.01) were grouped into clusters, and cluster significance was evaluated using the maximum cluster-sum statistic. Clusters surviving the corrected significance threshold (α = 0.025, two-tailed) were considered significant. Time window showing significant effects that were preserved across condition pairs were selected for further analysis (Gillis et al., 2021; Maris & Oostenveld, 2007).

Within each selected time window, we aimed to characterize the dominant TRF deflection. When a clearly defined local extremum was present, the dominant peak was identified as the local maximum (for positive components) or minimum (for negative components) within the window of interest. To improve the temporal resolution of peak latency estimates (approximately 8 ms at a sampling rate of 128 Hz), a parabola was fitted through the extremum and its two neighbouring samples, and peak amplitude and latency were defined as the amplitude and timing of the extremum of the fitted parabola.

In cases where a clearly defined local extremum could not be reliably identified across all subjects and conditions, peak responses were additionally quantified using a window-based extrema approach. Specifically, peak amplitude and latency were defined as the maximum (for positive components) or minimum (for negative components) TRF value within the predefined analysis window, without imposing constraints on local peak morphology. The search windows were defined as between 0 and 100 ms for P1, 50 and 150 ms for N1, 100 and 250 ms for P2 and 250 to 400 ms for N2. These ranges were chosen based on previous internal datasets (Dieudonné et al., 2024) as well as time windows previously found and used in the literature in continuous speech tasks (Borges et al., 2025; Ihara et al., 2021; Karunathilake et al., 2023). This proxy measure was used to enable consistent quantification across subjects and conditions and was treated as complementary to the dominant-peak analysis.

Finally, we examined the spatial distribution of the TRF weights. This analysis was carried out separately for each SNR. Within each pre-defined given window and SNR, we retained only the time points at which a cluster survived the permutation test in both conditions — that is, the intersection of the two surviving clusters within that window. TRF weights were then averaged across these retained time points, and the resulting scalp distributions were compared between the separated and collocated conditions.

### Statistical analysis

Signal processing and neural tracking calculations were performed in MATLAB (version 2025b, The Mathworks Inc., 2025). Cluster permutation test for analysis of temporal and spatial effects in the TRFs were performed using FieldTrip (Oostenveld et al., 2011). All the other statistical analysis was performed in R.

Neural tracking of speech has previously been used to estimate speech reception thresholds (SRTs) by modelling it as a sigmoid function of SNR and identifying its midpoint (Vanthornhout et al., 2018). Unlike linear models, the sigmoid provides interpretable parameters — a midpoint, a slope, and a location-dependent midpoint shift — the last of which yields a direct estimate of the spatial release from masking in decibels, in the same units as the behavioural SRM. Because spatial release from masking would manifest as a horizontal shift of this sigmoid along the SNR axis — equivalent to a shift in SRT — we fitted a non-linear mixed-effects model in which the midpoint of the sigmoid was allowed to vary with listening condition, using the *nlme* package (Pinheiro et al., 2025):

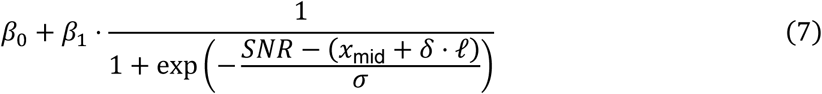

where *β*_0_is a baseline offset, *β*_0_ + *β*_1_is the asymptotic amplitude, *x*_mid_ is the SNR midpoint in the reference condition, *δ*quantifies the shift in midpoint per unit of location effect, ℓ is the effect-coded location regressor (ℓ = 1 for collocated, ℓ = −1 for spatially separated), and *σ* controls the steepness of the function. The parameters *β*_0_, *β*_1_, *x*_mid_, and *δ* were modelled as fixed effects. Under this coding, the sigmoid midpoint is *x*_mid_ + *δ* in the collocated condition and *x*_mid_ − *δ* in the spatially separated condition, such that spatial release from masking corresponds to a horizontal shift of 2*δ* along the SNR axis.

The model was fitted separately for neural tracking and behavioural self-reported speech intelligibility scores. For behavioural scores, which are bounded between 0 and 100%, *β*_0_ and *β*_1_were fixed to 0 and 100% respectively, and a by-subject random effect was placed on *x*_mid_ to capture individual differences in SRT. This specification is appropriate because individual differences in the asymptotic amplitude are constrained by the measurement scale, leaving the midpoint as the primary locus of between-subject variability. For neural tracking, *β*_1_ reflects the overall magnitude of cortical speech tracking, which varies substantially across participants due to individual differences in signal quality and neural responsiveness; a by-subject random effect on *β*_1_was therefore used instead. In both cases, the chosen random effect structure was verified to provide the best fit to the data by comparing alternative specifications.

To validate that the sigmoid model provided a better characterisation of the data than a simpler linear relationship, we compared it against linear mixed-effects models fit to the same data. For pairwise comparisons we used Two-sided Wilcoxon sign-rank tests as implemented in the *rstatix* package (Kassambara, 2023). When necessary, multiple comparison correction was performed through the Benjamini-Hochberg procedure (Benjamini & Hochberg, 1995). Effect sizes were calculated as the z-scores divided by the number of observations (Field et al., 2012). Effect sizes greater than 0.5 indicate a large effect (Cohen, 1992).

## Results

### Adaptive SRT estimation

We determined speech reception thresholds for each masker location with the Matrix sentence test through a behavioural adaptive procedure. All values are reported as average ± 1 standard deviation (s.d.). Individual SRTs are shown in **Figure 1**. As expected, subjects showed a decrease in speech reception threshold between the collocated and separated conditions. Average SRT for the collocated condition was −9.3 ± 0.9 dB SNR, while in the separated condition it was −15.7 ± 1.5 dB SNR. All subjects showed a decrease in SRT after separating target and masker (Wilcoxon sign-rank test p < 0.001, effect size = 0.877).

**Figure 1.**
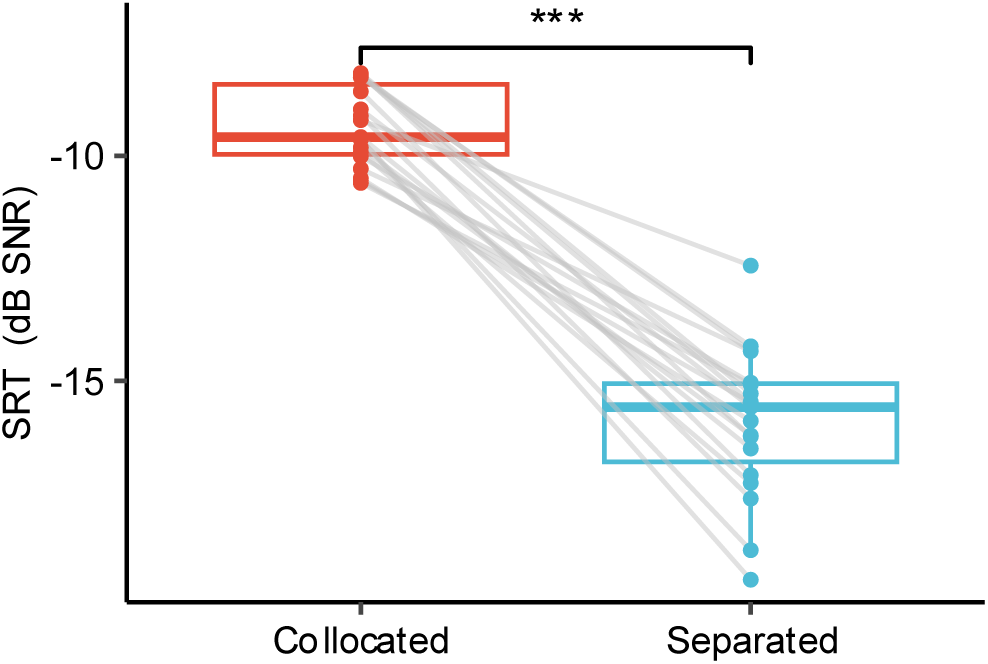
SRTs per subject for the collocated and separated conditions obtained with the Matrix sentence test (Two-sided Wilcoxon sign-rank test; ***p<0.001).

Average spatial release from masking was determined as the average difference between the separated and collocated condition across subjects and was 6.7 ± 1.7 dB.

We also analysed the effect of masker position (Left vs Right) in our setup. Results showed a smaller release from masking when the masker was in the right (+90° azimuth; mean SRM = 5.6 ± 1.5 dB) compared to when the masker was positioned in the left (−90° azimuth; mean SRM = 7.8 ± 1.3 dB). However, differences were not significant between corresponding conditions (Wilcoxon rank-sum test, p=0.079 for collocated, p=0.065 for separated). Although SRT in corresponding configurations were not significantly different, the release from masking measured in each subject that had the configuration S0N90 was lower than subjects that listened to S0N−90 (Wilcoxon rank-sum test, p < 0.01, effect size = 0.618).

### Self-reported speech understanding

We obtained self-reported percentage of speech understanding for each of the story fragments presented in trials with constant SNRs. To capture the bounded, saturating relationship between speech intelligibility and SNR, we fitted a logistic nonlinear mixed-effects model in which self-reported percentage of understood words followed a sigmoid function of SNR with a location-dependent midpoint. This model provided a substantially better fit than a linear mixed-effects model with the same predictors (ΔAIC = 151.5) modelled as *∼SNR*location + (1|subject).* The estimated midpoint in the reference condition was *x*_mid_= −8.6 dB SNR, and the location-dependent shift was *δ*= −2.6 dB SNR, such that the midpoint was −11.2 dB SNR in the spatially separated condition and −6.0 dB SNR in the collocated condition. Spatial release from masking thus corresponded to a shift of 2*δ*= 5.2 dB SNR along the SNR axis. The location shift was significant (*t*(205) = −30.82, p < 0.001), as was the steepness parameter (*σ* = 1.93 dB, *t*(205) = 24.22, p < 0.001). Pairwise comparisons on self-reported intelligibility, shown in **Figure 2A**, showed that intelligibility was significantly higher in the separated condition at every SNR tested (−13 to −4 dB: p < 0.001; −1 dB: p = 0.002), with consistently large effect sizes (≥ 0.84), strongest at the lowest SNRs (−13, −10, and −8 dB).

**Figure 2.**
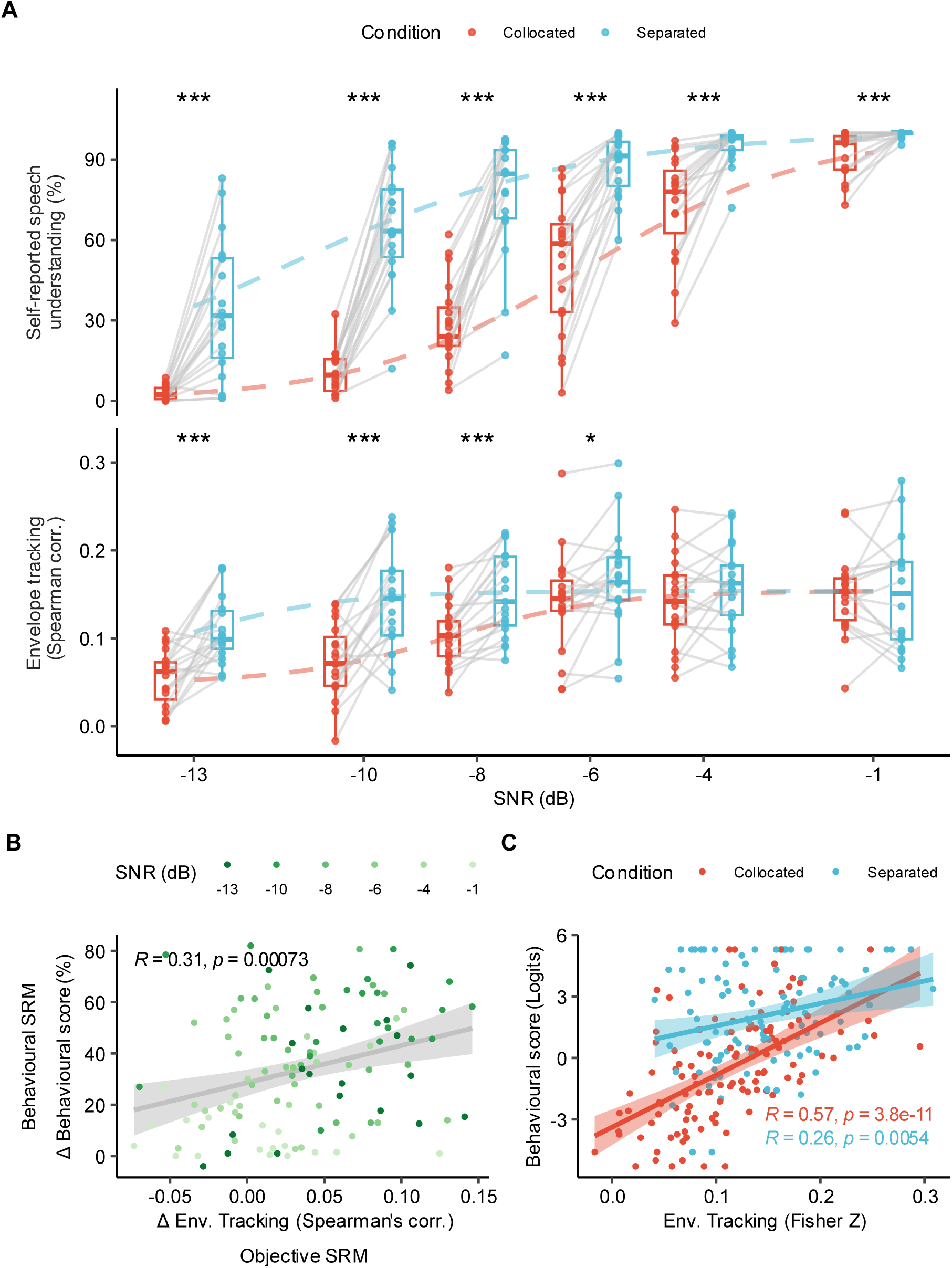
**A.** Behavioural scores and envelope tracking for each SNR and spatial condition. Points show individual-subject averages, and dashed lines indicate the predictive trends from a sigmoidal nonlinear mixed-effects model. (*p < 0.05, **p < 0.01, ***p < 0.001; two-sided Wilcoxon signed-rank test, FDR corrected).**B.** Differences between separated and collocated conditions for both untransformed measures, shown per SNR and subject. R and p-values correspond to Pearson’s correlation. **C.** Relationship between behavioural scores and envelope tracking for collocated and separated conditions. Behavioural scores were logit-transformed to reduce ceiling effects, and envelope tracking (Spearman correlations) was normalised using a Fisher Z transform.

### Backward model results

Envelope tracking was obtained for each of the 12 SNR x spatial-location combinations, leading to 226 values (2 conditions × 6 SNRs × 19 subjects; one subject contributed 10 rather than 12 values due to a missing SNR). Significant envelope tracking was defined as correlation values exceeding a 95% significance threshold derived from a permutation-based null distribution. This null distribution was obtained by correlating the original pre-processed speech envelope with circularly time-shifted versions of itself, thereby preserving the temporal structure and autocorrelation of the signals, estimated from 1,000 resamplings. This procedure was performed independently for each of the 12 combinations within each subject. Only 5 out of the 226 neural tracking values fell below this threshold. All five were in the collocated condition, at −13 dB SNR (n = 2) and −10 dB SNR (n = 3).

We modelled neural envelope tracking with the same logistic nonlinear mixed-effects model introduced above, with tracking following a sigmoid function of SNR with a location-dependent midpoint. This model provided a substantially better fit to the tracking data than a linear mixed-effects model with the same predictors (ΔAIC = 50.4, modelled as *∼SNR*location + (1|subject)*). The fitted group-level sigmoid functions for both neural tracking and self-reported scores are shown in **Figure 2A**.

The sigmoid model provided a good fit to the neural tracking data. The estimated SNR midpoint in the reference condition was *x*_mid_= −10.9 dB SNR, and the location-dependent shift was *δ*= −2.4 dB SNR, such that the sigmoid midpoint was −13.3 dB SNR in the spatially separated condition and −8.5 dB SNR in the collocated condition. Spatial release from masking thus corresponded to a shift of 2*δ*= 4.8 dB SNR along the SNR axis. The location shift was significant (*t*(203) = −10.17, p < 0.001), as were the baseline offset (*β*_0_ = 0.049, *t*(203) = 5.31, p < 0.001), asymptotic amplitude (*β*_1_ = 0.105, *t*(203) = 7.19, p < 0.001), and steepness parameter (*σ* = 1.40 dB, *t*(203) = 4.14, p < 0.001).

The horizontal shift between the two sigmoid curves implies that the benefit of spatial separation is largest in the SNR range bracketed by the two midpoints (−13.3 to −8.5 dB) and diminishes at more extreme SNRs where both conditions either saturate or fall to baseline. To examine this SNR-specific pattern directly, we conducted pairwise comparisons between conditions at each SNR level. Significantly higher envelope tracking was observed in the Separated than for the Collocated conditions at SNRs of −13, −10, −8, and −6 dB. The largest effects occurred at −13, −10, and −8 dB (effect sizes = 0.80, 0.77, and 0.87, respectively), while the effect at −6 dB was moderate (0.55). No significant differences were observed at −4 or −1 dB. **Table 1** contains the detailed results for this comparison. Importantly the increase between Collocated and Separated was consistent in almost all subjects for −8 dB (n=18 out of 19) and −10 dB (n=17 out of 19). **Figure 2A** also shows the pairwise comparison between spatial conditions across SNRs.

**Table 1.** Statistical values for pairwise comparison between envelope tracking values for separated and collocated conditions per SNR (Two-sided Wilcoxon sign-rank test).

| SNR (dB) | N-pairs | Statistic (W) | p-value | Effect Size | p-value (FDR corrected) |
| --- | --- | --- | --- | --- | --- |
| -13 | 19 | 8 | $<10^{-4}$ | 0.803 | $<0.001$ |
| -10 | 19 | 12 | $<0.001$ | 0.766 | $<0.001$ |
| -8 | 19 | 1 | $<10^{-5}$ | 0.868 | $<10^{-4}$ |
| -6 | 19 | 36 | 0.016 | 0.545 | 0.024 |
| -4 | 19 | 63 | 0.210 | 0.295 | 0.252 |
| -1 | 18 | 92 | 0.799 | 0.067 | 0.799 |

**Table 2.** Statistical values for pairwise comparison for P2 latencies and amplitudes for separated and collocated conditions per SNR (Two-sided Wilcoxon sign-rank test, FDR corrected).

| SNR (dB) | N-pairs | Statistic (W) | p-value | Effect Size | p-value (FDR corrected) |
| --- | --- | --- | --- | --- | --- |
| Latencies |  |  |  |  |  |
| -13 | 19 | 88 | 0.798 | 0.066 | 0.798 |
| -10 | 19 | 153 | 0.018 | 0.535 | 0.027 |
| -8 | 19 | 164 | 0.004 | 0.637 | 0.016 |
| -6 | 19 | 159 | 0.008 | 0.591 | 0.016 |
| -4 | 19 | 159 | 0.008 | 0.591 | 0.016 |
| -1 | 18 | 114 | 0.229 | 0.293 | 0.275 |
| Amplitudes |  |  |  |  |  |
| -13 | 19 | 61 | 0.182 | 0.314 | 0.338 |
| -10 | 19 | 64 | 0.225 | 0.286 | 0.338 |
| -8 | 19 | 6 | $<10^{-4}$ | 0.822 | $<0.001$ |
| -6 | 19 | 85 | 0.709 | 0.092 | 0.734 |
| -4 | 19 | 46 | 0.049 | 0.452 | 0.148 |
| -1 | 18 | 94 | 0.734 | 0.087 | 0.734 |

### Relationship between behavioural and neural measures

We examined the relationship between behavioural performance and neural envelope tracking in two ways: first using the spatial-separation benefit (the difference between the separated and collocated conditions) and second using the absolute scores within each condition. The behavioural benefit, quantified as the percentage difference in self-reported word recognition between spatially separated and collocated conditions for each SNR and subject, showed a significant positive linear relationship with the corresponding increase in neural tracking (Pearson’s r = 0.31, t(105) = 3.29, p < 0.001). This relationship is illustrated in **Figure 2B**.

We also examined the direct relationship between the behavioural scores and neural tracking, rather than the spatial-separation benefit. Both measures are bounded variables whose raw distributions are compressed near their limits, violating the assumptions of linear analysis; both were therefore transformed before examining their relationship. The self-reported scores are percentages bounded between 0 and 100 and showed ceiling and floor effects, so they were logit-transformed. The neural-tracking values are Spearman correlation coefficients, whose sampling distribution is skewed and bounded at ±1; they were therefore Fisher-transformed to place them on an unbounded, variance-stabilised scale. Behavioural scores were positively correlated with neural tracking in both conditions, with a stronger relationship in the collocated condition (Pearson’s r = 0.57, p < 0.001) than in the separated condition (Pearson’s r = 0.26, p = 0.005). These correlations are shown in **Figure 2C**. Because ceiling effects were still found after the transformations, we also calculated the correlation between all the obtained behavioural and neural scores after ignoring results for SNR = −1 dB (Pearson’s r = 0.54, p < 0.001). This can be seen in **Supplementary Figure 1**.

### TRF results

Cluster-based permutation testing revealed significant TRF response clusters across conditions, with the most consistent effects across SNRs occurring between approximately 150 ms and 250 ms. These clusters motivated the focus on this latency range, which corresponds to the standard mid-latency TRF components. **Figure 3A** shows the average shape of the TRFs across subjects for each tested condition as well as the latencies in which the cluster permutation test showed a consistent deflection. In addition, increasing SNR was associated with more consistent responses both at early latencies following stimulus onset (40–60 ms) and at later latencies around 300 ms.

**Figure 3.**
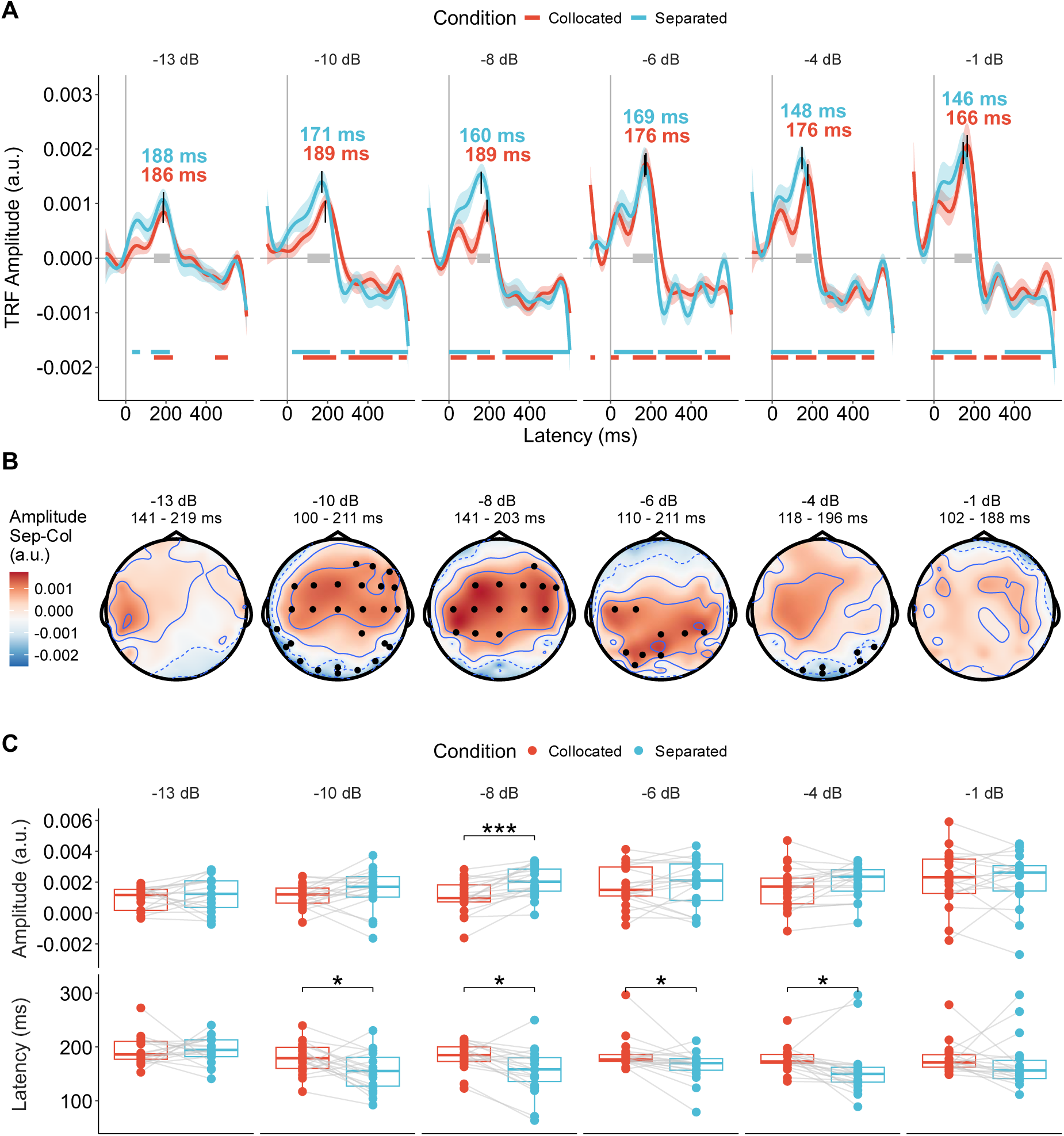
**A.** Average TRFs across subjects for each SNR and spatial condition. Shaded ribbons indicate the standard error of the mean. The marked peak corresponds to P2, with its latency (ms) indicated separately for the collocated and separated conditions. Coloured horizontal bars indicate time windows where the TRF amplitude differed reliably from zero across subjects (cluster-permutation test); grey bars mark the overlap between conditions, identifying windows where both retained a significant cluster around the P2 peak. These windows were used to define the time regions for the spatial analysis shown in the bottom panel. **B.** Topographic amplitude differences of the TRF averaged within the P2 window (grey segment in Top). Markers indicate clusters that remained significant after cluster-permutation correction. **C.** Subject pairwise comparison of P2 latencies across conditions (*p < 0.05, **p < 0.01, ***p < 0.001; two-sided Wilcoxon signed-rank test, FDR corrected).

Peak amplitude and latency were operationally defined as the maximum (for positive components) or minimum (for negative components) TRF value within predefined time windows of interest corresponding to the standard components P1, N1, P2, and N2 (see Methods). These window-based extrema were extracted for each subject and condition and entered into linear mixed-effects models with SNR and spatial condition (collocated vs. separated) as fixed effects and subject as a random intercept, explicitly acknowledging that for some components the extracted value may not reflect a clearly identifiable physiological peak. SNR = −1 dB was excluded to include all subjects (n = 19).

All four components showed significant amplitude growth with SNR (P1: p < 0.001; N1: p = 0.018; P2: p < 0.001; N2: p < 0.001). P2 and N2 latencies decreased significantly with SNR (P2: p = 0.002; N2: p = 0.030), while P1 and N1 latencies were SNR-invariant. Regarding spatial condition, P1 showed greater amplitude (p = 0.017) and later latency (p = 0.007) in the separated condition, and P2 peaked earlier when noise was separated (p < 0.001). N1 and N2 were insensitive to spatial condition in both amplitude and latency. A significant SNR × location interaction was found only for P2 amplitude (p = 0.040), indicating attenuated SNR-dependent amplitude growth in the separated condition.

Post-hoc analyses were deliberately restricted to components exhibiting consistent morphology across subjects. Accordingly, pairwise comparisons were performed only for the P2 component, which was the only peak that could be reliably identified across all subjects and conditions. For the remaining components, peak identification was inconsistent, and the window-based extrema should be interpreted as descriptive rather than reflecting robust, physiologically distinct peaks. **Supplementary Figure 2** therefore reports amplitudes and latencies for P1, N1, P2, and N2 obtained using the window-based extrema approach, illustrating that only P2 exhibits consistent and interpretable trends across conditions. No further pairwise analyses were conducted for P1, N1, or N2.

Pairwise comparisons across SNRs for the P2 component revealed a significant decrease in latency with spatial separation for all conditions except −13 and −1 dB as shown in **Figure 3C**. With respect to amplitude, P2 peaks were significantly larger in the separated condition at −8 dB.

Finally, we examined the spatial distribution of TRF weights across the scalp. **Figure 3B** indicates the window across which the channel amplitudes were averaged which corresponded to the time lags in which both conditions showed a significant effect. Significant condition-dependent effects were observed at −10, −8, −6, and −4 dB. These effects were driven by a cluster of central electrodes showing increased amplitudes, accompanied by a decrease in amplitude over left-lateralized and posterior electrodes.

## Discussion

### Behavioural results and spatial release from masking

The present study confirmed that SRM is robustly observed in normal-hearing listeners. Participants showed decreased speech reception thresholds when target and masker were spatially separated, with an average SRM of 6.7 dB as measured in the adaptive procedure. These results also agree with the self-reported percentage of understood words for the stories: as expected, the largest benefits occurred at the lowest SNRs, highlighting the SNR-dependent nature of binaural unmasking. Subtle asymmetries between left- and right-side maskers may reflect potential right-ear advantage effects or small acoustic asymmetries in the experimental setup, consistent with prior behavioural findings (Garadat et al., 2009; Williges et al., 2015). Across all tested SNRs, self-reported speech understanding was consistently higher in the separated condition, with effect sizes strongest at the lowest SNRs, emphasising the functional importance of spatial cues in challenging listening conditions.

### Neural envelope tracking

Building on these behavioural observations, backward-model analyses show that spatial separation enhances cortical speech tracking, especially at low SNRs (−13 to −6 dB), with the SNR–tracking function shifted toward lower SNRs in the spatially separated condition. Notably, these effects were observed in 17 out of 19 subjects at intermediate low SNRs, suggesting the potential subject-specific reliability of neural tracking measures as indicators of spatial unmasking. These results suggest that neural SRM parallels behavioural SRM, highlighting the potential value of envelope tracking as an objective marker of spatial unmasking. In contrast, previous work by Song et al. (2020) reported no overall effect of spatial separation on envelope tracking. In their study, which investigated the effects of masker type, spatial location and language experience, the collocated and separated conditions were presented at different SNRs in order to equate behavioural performance, which may explain the absence of a spatial-separation effect. By testing across a broad SNR range (−13 to −1 dB) under energetic masking, we consistently observed neural benefits at challenging SNRs, revealing spatial unmasking effects that narrower designs may miss. These findings align with Dieudonné et al. (2024), who reported increased neural tracking linked to binaural release from masking, and with Ahmed et al. (2026) who found that adults show stronger tracking of target speech when competing talkers were spatially separated. Converging evidence comes from EEG-based auditory attention-detection paradigms, in which decoding accuracy calculated from envelope tracking improves as competing talkers are more separated in space (Das et al., 2018). Collectively, these studies suggest that spatial separation enhances neural tracking under adverse listening conditions, and that this neural benefit reflects the speech-understanding gains associated with spatial hearing.

### TRFs and cortical dynamics

TRF analyses revealed condition-dependent modulation of both timing and amplitude, with prominent deflections between 150–250 ms corresponding to canonical N1–P2–N2 components. P2 amplitude and latency emerged as particularly sensitive to spatial configuration and SNR, showing larger amplitudes and earlier latencies in separated, low-SNR conditions. At higher SNRs, the clusters in which the TRF differed from zero across subjects (i.e. the time windows surviving the permutation threshold) were more extensive in both spatial conditions. The relatively short recording durations per condition likely contributed to inter-subject variability and may have limited sensitivity to more subtle spatial effects, including the resolution of individual peaks; extending recording times and increasing the number of trials would improve reliability and allow for better separation of early sensory versus later integrative components (Crosse et al., 2021).

Spatial distribution analyses demonstrated central electrode activity increases with spatial separation, consistent with engagement of broader cortical networks. Such redistribution parallels findings from ERP and ACC studies, indicating recruitment of additional cortical resources under complex listening scenarios. These results support the interpretation of TRFs as continuous-speech counterparts of classical AEPs, capturing hierarchical processing from early acoustic encoding to later integrative stages sensitive to spatial unmasking (Dieudonné et al., 2024; Kim & Epp, 2023; Li et al., 2024; Zobel et al., 2022). The observed TRF patterns align with literature on cortical auditory evoked potentials, where spatial separation increases P2 amplitudes, shortens latencies, and reflects perceptually relevant unmasking (Fan & Gifford, 2024; Papesh et al., 2017). TRFs extend this framework to continuous speech, with later components such as P2 showing sensitivity to speech intelligibility beyond purely acoustic representation. Binaural unmasking effects observed in TRFs, such as decreased latencies and increased P2 amplitudes, mirror findings from comparable neural-tracking and evoked-potential studies under both low SNR and masked conditions (Dieudonné et al., 2024; Kim & Epp, 2023; Zobel et al., 2022), suggesting that TRFs are sensitive to the same effects as those measures. Consistent with these TRF observations, earlier studies suggest that binaural unmasking arises primarily at cortical stages. Brainstem ASSRs show minimal BMLDs, whereas cortical ASSRs and FFRs reflect sensitivity to binaural cues and perceptual SRM (Clinard et al., 2016; Koerner et al., 2020; Wong & Stapells, 2004). The present results reinforce this hierarchy, as spatial separation enhanced cortical tracking selectively under adverse conditions rather than producing uniform gains across the auditory pathway, indicating the potential role of higher-level cortical mechanisms in exploiting spatial cues. It should be noted, however, that brainstem responses are generally harder to record than cortical responses (Bidelman et al., 2018; Maddox & Lee, 2018); the observation that binaural-unmasking effects emerge mainly at the cortical level may therefore stem in part from this difference in measurement sensitivity rather than from a true cortical origin of the effects.

### Implications for difficult-to-test populations

The convergence of behavioural and neural SRM supports the use of neural tracking as a flexible and objective framework for assessing spatial hearing. An objective measure is especially valuable for populations that are difficult to test behaviourally, such as young children or listeners who cannot provide reliable responses, where neural tracking can probe how auditory pathways transmit spatial cues without requiring an overt behavioural task. By establishing normative patterns in normal-hearing listeners, these findings provide benchmarks for evaluating both cortical processing and binaural benefits in clinical and device-based interventions (Balkenhol et al., 2020; Dieudonné et al., 2024; Gillis et al., 2022; Sasaki et al., 2009). Building on the observed correspondence between behavioural and neural SRM, beyond TRFs, neural tracking metrics could be used to assess continuous speech processing under varying SNRs and spatial configurations, capturing unmasking, redundancy, and squelch effects, and may serve as a useful tool for evaluating binaural listening and informing hearing-device assessments. Employing more diverse acoustic scenarios and SNRs could further elucidate how cortical speech tracking adapts to complex and realistic listening environments, refining our understanding of the neural mechanisms underlying spatial unmasking and SRM.

## Conclusion

In summary, the study demonstrates that spatial separation robustly enhances cortical speech tracking and closely mirrors behavioural SRM. TRFs serve as continuous-speech analogon of AEPs, capturing hierarchical auditory processing influenced by SNR and spatial configuration. These findings support neural tracking as a sensitive and objective measure of spatial hearing and provide a foundation for future work in clinical populations and hearing-device research, offering both mechanistic insights and practical avenues for assessment and intervention.

## Acknowledgements

This work is part of Horizon-MSCA-2022-DN-01: CherISH; a European Doctorate Network project funded by the European Union’s Horizon 2020 framework program for research and innovation under the Marie Sklodowska-Curie Grant Agreement No: 101120054.

## Supplementary figures

**Supplementary Figure 1.**
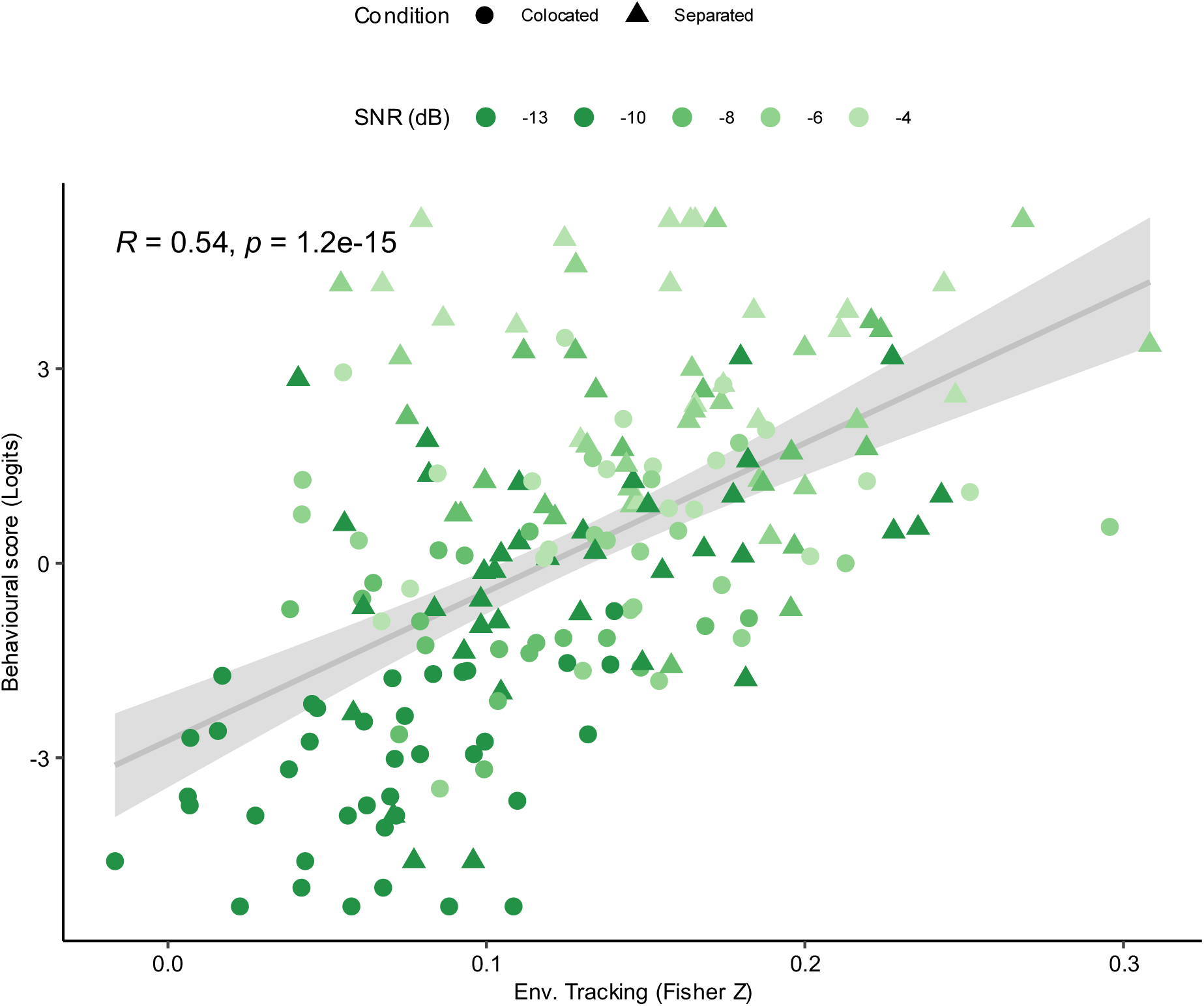
Average scores per SNR and location (Logit transformed) vs envelope tracking (Fisher Z transformed) for each subject. SNR=-1 dB was removed in this figure as ceiling effects were present in both conditions for the behavioural score across all subjects. Different shapes indicate collocated or separated conditions. Pearson’s R corresponds to the linear correlation between behavioural and envelope tracking scores. P-value corresponds to the t-statistic associated with the R value.

**Supplementary Figure 2.**
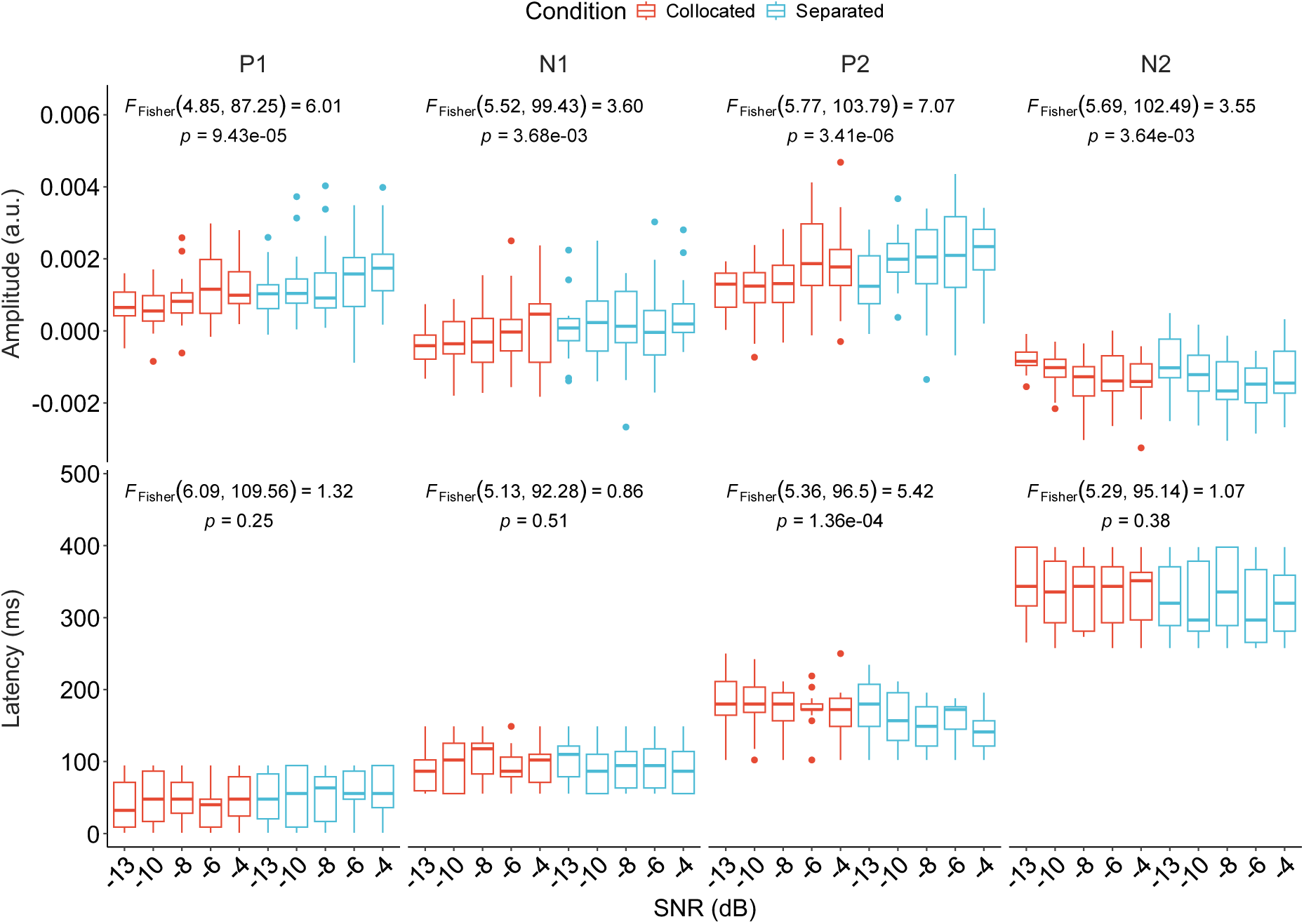
Amplitude and latency of the first two negative and positive peaks of the TRFs across SNR and location. Peaks were defined as the maximum or minimum amplitude within predefined time windows of interest. Statistical annotations show results from repeated-measures ANOVAs, included for descriptive purposes. Full inferential analyses using linear mixed-effects models are reported in the main text.

